# Cooperativity in septin polymerization is tunable by ionic strength and membrane adsorption

**DOI:** 10.1101/2025.02.12.637902

**Authors:** Ellysa Vogt, Ian Seim, Wilton T. Snead, Amy S. Gladfelter

**Affiliations:** Curriculum in Genetics and Molecular Biology, University of North Carolina at Chapel Hill, 27514, Chapel Hill, North Carolina, United States; Department of Cell Biology, Duke University Medical School, 27710, Durham, North Carolina, United States; Max Planck Institute of Molecular Cell Biology and Genetics, Pfotenhauerstraße 108, 01307 Dresden, Germany

**Author notes:** Department of Cell and Developmental Biology, Feinberg School of Medicine, Northwestern University, 60611, Chicago, Illinois, United States (Wilton T. Snead).

## Abstract

Cells employ cytoskeletal polymers to move, divide, and pass information inside and outside of the cell. Previous work on eukaryotic cytoskeletal elements such as actin, microtubules, and intermediate filaments investigating the mechanisms of polymerization have been critical to understand how cells control the assembly of the cytoskeleton. Most biophysical analyses have considered cooperative versus isodesmic modes of polymerization; this framework is useful for specifying functions of regulatory proteins that control nucleation and understanding how cells regulate elongation in time and space. The septins are considered a fourth component of the eukaryotic cytoskeleton, but they are poorly understood in many ways despite their conserved roles in membrane dynamics, cytokinesis, and cell shape, and in their links to a myriad of human diseases. Because septin function is intimately linked to their assembled state, we set out to investigate the mechanisms by which septin polymers elongate under different conditions. We used simulations, *in vitro* reconstitution of purified septin complexes, and quantitative microscopy to directly interrogate septin polymerization behaviors in solution and on synthetic lipid bilayers of different geometries. We first used reactive Brownian dynamics simulations to determine if the presence of a membrane induces cooperativity to septin polymerization. We then used fluorescence correlation spectroscopy (FCS) to assess septins’ ability to form filaments in solution at different salt conditions. Finally, we investigated septin membrane adsorption and polymerization on planar and curved supported lipid bilayers. Septins clearly show signs of salt-dependent cooperative assembly in solution, but cooperativity is limited by binding a membrane. Thus, septin assembly is dramatically influenced by extrinsic conditions and substrate properties and can show properties of both isodesmic and cooperative polymers. This versatility in assembly modes may explain the extensive array of assembly types, functions, and subcellular locations in which septins act.

**SIGNIFICANCE:** The septin cytoskeleton plays conserved and essential roles in cell division, membrane remodeling, and intracellular signaling with links to varied human diseases. Unlike actin and microtubules, whose polymerization dynamics have been extensively characterized, the molecular details of septin polymerization remain poorly understood. Here, we investigate the mode of septin polymerization through the lens of isodesmic and cooperative polymer assembly models in solution, on planar and curved supported membranes, and under different ionic conditions. Our findings show that the mechanisms of septin assembly are highly sensitive to ionic conditions, membrane geometry, and protein concentrations. Notably, assembly can show either cooperative or isodesmic properties depending on context, thereby revealing unexpected plasticity.

## INTRODUCTION

Cytoskeletal proteins such as actin and microtubules are present in a soluble state in all cells and their function emerges through highly regulated polymerization. Cells control the location, time, length, and geometry of these polymers with precision. Integral to this regulation are intrinsic properties of the polymers themselves, such as dynamic instability or treadmilling that are encoded in the structural and enzymatic features of the proteins. Biophysical descriptions of the polymerization process have been critical for understanding how the intrinsic features of the proteins are harnessed for regulation. The modes of polymerization are integral to the ability of cells to control polymer length, concentration-dependent rates of formation, and impact the types of regulatory proteins that are employed to tune these properties (*1*). For example, actin cables are built in polarized cells to target secretion to specific sites using nucleators that facilitate cooperative assembly of F-actin. Thus, describing the fundamental properties of polymerization has been critical to understanding both the form and function of the cytoskeleton in cells.

The septin cytoskeleton shares some basic properties with each of the more well-studied cytoskeletal components. Like microtubules and actin, septins can bind and hydrolyze nucleotides, unlike intermediate filaments that assemble without complexing or hydrolyzing a nucleotide (*2-4*). However, unlike microtubules and actin but similar to intermediate filaments, septin filaments assemble with no known structural or functional polarity (*5*). Nevertheless, certain septin assemblies in cells, for example at the mother-bud neck in budding yeast, display apparent asymmetries of unknown origins.

The basic subunits of septin polymers are hetero-oligomeric complexes that are co-translationally assembled octamers (*6*). Each octamer consists of two copies of 4 different proteins linearly arranged in a palindromic order, creating a rod that is 32 nm long. These hetero-oligomers are thought to act as the soluble, protomeric form of septins in the cytosol, which then elongates when bound to membranes or other cytoskeletal elements (**Fig 1A**) (*7-9*). Recombinant septins can also polymerize in solution at sufficiently high concentrations and in a salt-dependent manner (**Fig 1A**) (*10-12*). Association with diffusive lipid membranes can promote septin polymerization in a membrane curvature-dependent manner, with polymerization occurring most rapidly on micron-scale curved membranes. As such, septins are the first described sensors of micron-scale membrane curvature in eukaryotes. Interestingly, preferential assembly on specific membrane geometries is kinetically-driven, highlighting the remarkable ability of these nanometer-scale complexes to discern shallow curvatures from flat membranes that should seemingly appear identical (*2, 5, 9, 11, 13-16*). Despite this basic understanding of key features of assembly, the mode of septin polymerization has not been extensively studied. The goal of this study is to characterize the mechanism of septin assembly *in vitro* so as to understand how cells may manipulate these fundamental properties to control assembly in time and space.

**Figure 1:**
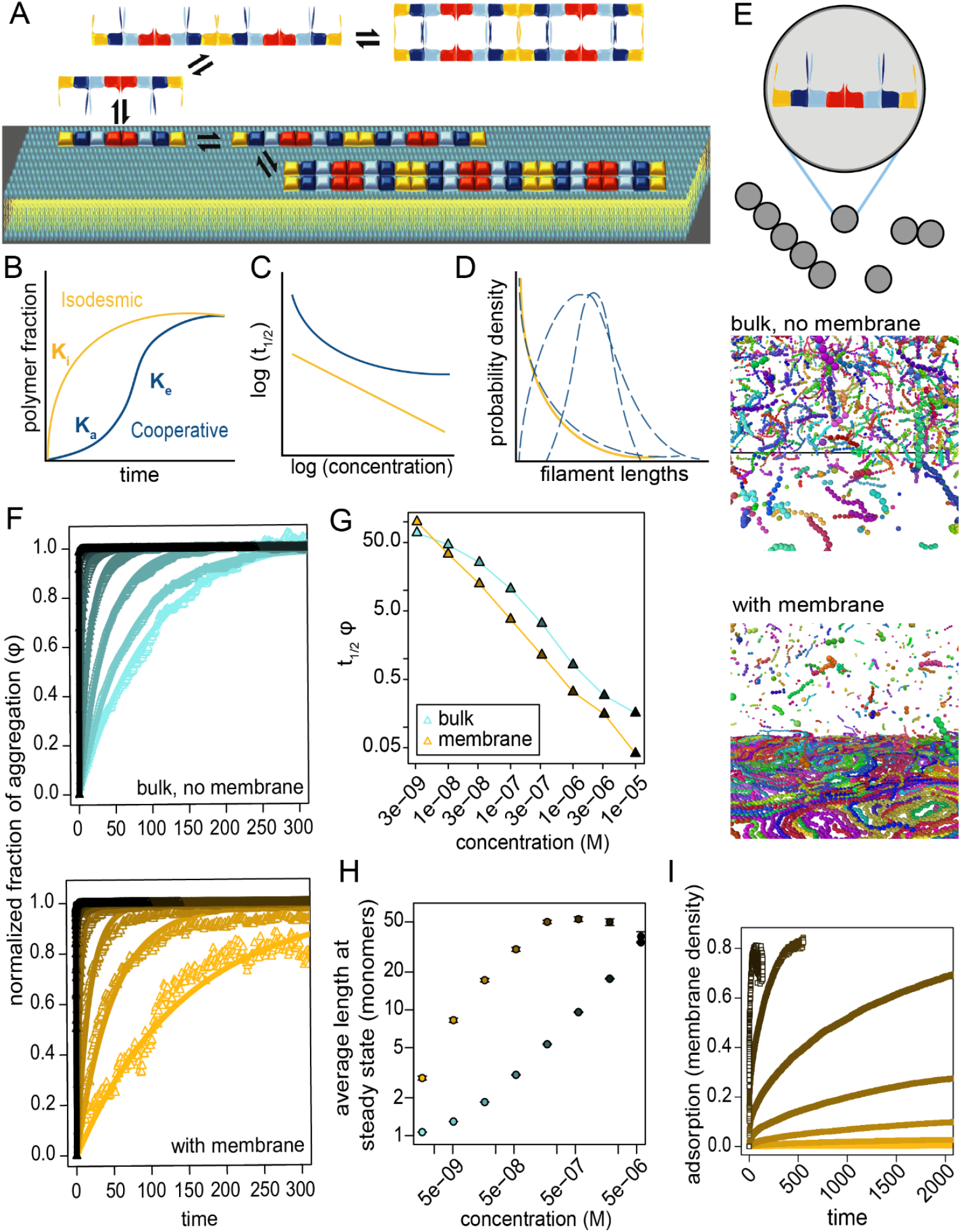
Membrane binding is not sufficient to induce cooperative assembly. A) Cartoon diagram of a septin octamer and filament assembly reactions in solution and at a membrane. B) Predicted polymerization of cooperative and isodesmic polymers over time. C) Predicted log *t*_1/2_ as a function of concentration for cooperative and isodesmic polymers. D) Predicted filament length distributions of cooperative and isodesmic polymers at steady state. Cooperative polymers can exhibit multiple filament length distributions and are showed as dashed lines, while isodesmic polymers will only exhibit exponential curves. E) Top, In the simulation, octamers are represented as circles which diffuse freely and can bind and unbind other octamers and the membrane. Bottom, snapshots of steady-states in simulations of isodesmic polymers in bulk or in bulk with an attractive surface mimicking a membrane. F) Fraction of aggregation over time at multiple concentrations for simulations without a membrane (top) and with a membrane (bottom). G) log *t*_1/2_ for fraction of aggregation (*φ*) as a function of concentration for simulations without membranes (teal) and with a membrane (yellow). H) Mean filament lengths at steady state across a range of concentrations for simulations with membranes (yellow) and without membranes (teal). I) Simulated adsorption to the membrane over time.

Classic descriptions of polymer elongation mechanisms often describe cooperative or isodesmic assembly processes. These different modes of assembly display some distinct, measurable features that can distinguish them, including the rate of assembly, the concentration-dependent time to reach steady-state, and filament length distributions. To form stable assemblies, some cooperative polymers must overcome an energetically unfavorable nucleation step, resulting in a kinetic lag during assembly (**Fig 1B**) and a critical concentration below which polymerization is minimal (*17, 18*).

Once nucleation occurs, elongation is energetically favorable, and assemblies rapidly form. Nucleation can be achieved through a variety of mechanisms, including lateral interactions that stabilize monomers (*19*), conformational changes of subunits driven by bound nucleotide state (*17*), and the formation of helical filaments in which interactions among additional subunits and existing filaments become available after formation of the first loop (*18*). Because not all cooperative polymers exhibit unfavorable nucleation steps at all concentrations (*18*), we define cooperative polymerization in this study as the presence of at least two different association constants during assembly, implying length-dependent rates of polymerization and/or depolymerization (*1, 18*). We are interested in positive cooperativity, in which the association constant for small sizes, *K*_*a*_, is smaller than that of large sizes, *K*_*e*_, resulting in an increasing slope of time-dependent polymerization at early times (**Fig 1B**). A kinetic lag is only present at low concentrations when nucleation is energetically unfavorable, whereas for sufficiently high concentrations when nucleation is energetically favorable, immediate, albeit slower, growth can be observed at early times (*18, 20*). The ability to distinguish polymerization rates at early and late times in kinetic curves depends on the extent to which *K*_*a*_ < *K*_*e*_. Thus, several different parameters can vary, including the favorability of a nucleation step, the free concentration, and dissociation rates, but a key overall property of cooperative assemblies is the presence of two distinct association constants.

In contrast, non-cooperative or isodesmic polymerization can be described by length-independent rates of polymerization and de-polymerization, such that a single association constant, *K*_*i*_, characterizes the system. Without a weaker association constant at small sizes, isodesmic polymerization is most rapid when polymerization begins and steadily plateaus over time (**Fig 1B**). These kinetic signatures of cooperative versus isodesmic polymerization can also be distinguished in plots of *t*_1/2_, the time required for a system to reach half of its steady-state value, as a function of concentration (*20, 21*). For isodesmic polymers, *t*_1/2_ steadily decreases with concentration, while for cooperative polymers, *t*_1/2_ saturates at a minimum value and eventually becomes independent of concentration at high concentrations (**Fig 1C**).

Steady-state length distributions also provide some information about the underlying assembly dynamics of biological polymers (*1*). The length-independent assembly and disassembly rates characteristic of isodesmic systems imply that steady-state length distributions will always be exponential and therefore have no preferred length (**Fig 1D**) (*1*). In contrast, because cooperative systems have length-dependent assembly and/or disassembly rates, steady-state length distributions can be more varied. For example, in the case of a filament with an unfavorable nucleation step, a fast elongation rate, and a finite monomer sub-pool that is depleted at long times (*1*), the steady-state length distribution will have one peak corresponding to monomers at the critical concentration and a second peak at an average filament size (**Fig 1D**). However, some cooperative polymers can also lead to near-exponential or monotonically decreasing length distributions. For example, a polymer with an unfavorable nucleation step and a single elongation rate results in an exponential length distribution for sizes larger than the nucleus, which can be difficult to distinguish experimentally from a true exponential distribution when the nucleus size is small (**Fig 1D**, dashed lines) (*22*). With a fragmentation reaction added, length distributions are monotonically decreasing (*22*). Therefore, experimental observation of peaked steady-state length distributions always indicates that the system is not isodesmic, but observation of exponential or monotonically decreasing distributions cannot distinguish between isodesmic or cooperative dynamics.

By considering measurable properties of isodesmic versus cooperative polymers, we set out to characterize how septin filaments elongate under different conditions. We used coarse-grained simulations, *in vitro* reconstitution of purified septin complexes, and quantitative microscopy/spectroscopy to monitor septin polymerization behaviors on synthetic lipid bilayers and in solution under differing ionic strengths. We find that the mechanism of septin polymerization *in vitro* defies rigid categorization and instead shows a mixture of apparent isodesmic and cooperative assembly properties that are highly context-dependent. These data also support a significant electrostatic component in septin assembly and membrane adsorption. In much the same way that septins display a blend of properties typically associated with distinct classes of cytoskeletal filaments, including nucleotide use and lack of polarity, so too do septins exhibit a blend of polymerization mechanisms.

## METHODS

### Yeast septin protein purification

Recombinant S. cerevisiae septin complexes were expressed in BL21 (DE3) *Escherichia coli* cells transformed with a duet expression system (*9*). Selected transformants were grown overnight in 100 mL LB with ampicillin and chloramphenicol at 37º C. Four liters of terrific broth with selection antibiotics were inoculated with 32 mL of overnight culture and grown to an O.D. 600 nm between 0.6 and 0.8. Once at the desired density, cells were induced with 1 mM isopropyl-B-D-1-thiogalactopyranoside (IPTG) and grown for 18 h at 18º C. Pellets were harvested by centrifugation at 9,000 RCF for 15 min then frozen at -80º C. All further steps to process the cell pellets were done on ice or at 4º C.

Pellets were thawed in cold lysis buffer (1 M KCl, 50 mM HEPES pH 7.4, 10 mM imidazole, 1 mM MgCl2, 10% glycerol, 1% Tween-20), 1 mg/mL lysozyme, and 4 1x EDTA free protease inhibitor tablet (Pierce, 1 tablet per liter of culture). To generate cell lysates, the pellets were first resuspended with a 25 mL serological pipette until thin, homogenized with a 40 mL glass douncer, then sonicated on ice for 5 s on, 10 s off for 5 min. Lysate was clarified by centrifugation at 20,000 RPM for 30 min. Clarified lysate was incubated with 4 mL equilibrated cobalt resin for 1 h then poured over a gravity flow column. Once the resin was settled, wash buffer (1 M KCl, 50 mM HEPES pH 7.4, 20 mM imidazole) was flowed over the column 3x (12 mL first, then 5 mL for the second and third washes). Bound protein was eluted (300 mM KCl, 50 mM HEPES pH 7.4, 500 mM imidazole) then dialyzed into septin storage buffer (300 mM KCl, 50 mM HEPES pH 7.4, 10 % glycerol, 1mM BME).

After dialysis, the protein was incubated overnight with 42 µM SNAP-488 and 4 mM DTT at 4º C. Excess label was washed out using Zeba spin desalting columns before snap freezing in single use aliquots.

### Lipid mix preparation

In a glass vial, chloroform stocks of lipids were mixed then dried using argon gas followed by an overnight incubation under negative vacuum pressure. For planar assays, the lipid composition comprised 75 mole percent dioleoylphosphatidylcholine (DOPC), 25 mole percent phosphatidylinositol (soy, PI), and 0.05 mole percent of rhodamine-phosphotitdyl-ethanolamine (Rh-PE). Bead assay lipid composition were similar with 73% DOPC and 2 mole percent biotinylated-phosphotitdyl-ethanolamine. The evaporated lipid films were hydrated using a supported lipid bilayer buffer (SLBB, 150 mM KCl, 20 mM HEPES pH 7.4, and 1 mM MgCl_2_) to a final concentration of 5 mM. The lipid films were vortex mixed 10 s then incubated for 5 min at 37º C, this process was repeated at least 5 times to fully hydrate. To form small unilamellar vesicles, the hydrated films were bath sonicated until the opaque mixture became transparent.

Single use aliquots (10-30 µL) were gently treated with argon gas before snap freezing and stored at -80º C for no more than 3 months.

### Preparation and visualization of planar supported lipid bilayers

Glass coverslips were treated with plasma (Pe25-JW; Plasma Etch) on high power for 5 minutes and a reaction chamber (*38*) was applied to the center of the coverslip. SUVs were diluted to 1 mM in SLBB with 1 mM CaCl2 and incubated for 20 min at 37º C. Excess lipids were disrupted and removed though a series of washes (with the pipette tip in the center of the well, never contacting the coverslip) first with 150 µL of SLBB then with 150 µL of reaction buffer (50 mM HEPES pH 7.4, 200 µg/mL β-casein, 1 mM BME, ± 33.3 mM KCl). In the last wash, all liquid was carefully removed and replaced with 75 µL reaction buffer.

The rhodamine in the bilayer was used to set the critical angle for total internal reflection fluorescence microscopy (TIRF) using a Nikon Ti2 E-TIRF system with a LUN-4 laser launch with solid state 405/488/561/640 nm lasers and Hamamtsu Fusion BT sCMOS detector. The critical angle was set for the 561 channel and approximated for the 488 channel. An image of the bare bilayer was taken in 561 and 488 to be used for background subtraction and record of bilayer quality. 25 µL of septins at the desired concentration (in SSB either containing 300 mM KCl or 200 mM KCl for experiments with final KCl concentrations of 75 and100 mM KCL or 50 mM, respectively) were mixed in the reaction buffer, then the critical angle of the 488 channel was quickly adjusted as needed and the acquisition was started.

### Preparation and visualization of bead supported lipid bilayers

Lipid bilayers formed on 1 µm silica microspheres were generated (*38*). After bath sonicating the SUVs and beads to minimize clumping, 50 nM SUVs were added to 1 µm silica beads at a total surface area of 440 mM2 and incubated at room temperature on an end-over-end rotator for 1 h. To wash away excess lipids, the beads were gently pelleted at 2300 RCF and washed with 200 µL pre-reaction buffer (33.3 mM KCl and 50 mM HEPES pH 7.4), repeated for a total of 4 washes. The beads were diluted by mixing 29 µL washed beads into 721 µL reaction buffer (50 mM HEPES pH 7.4, 200 µg/mL β-casein, 1mM BME, ± 33.3 mM KCl).

Reaction chambers were prepared similar to the planar system on a PEG and 15% Biotin-coated coverslip (*13, 14*). The chamber was washed with 200 µL reaction buffer (50 mM HEPES pH 7.4, 0.1% methylcellulose, 200 µg/mL β-casein, 1 mM BME, ± 33.3 mM KCl), then 50 µL of 1 mg/mL NeutrAvidin was left in the chamber to bind the PEG-Biotin for at least 15 min. The NeutrAvidin was removed and 75 µL of diluted beads were added and allowed to settle and bind the NeutrAvidin link for at least 30 min.

Beads were imaged using a Zeiss 980 laser scanning confocal with Airyscan sensor. A single focal plane in the middle of the beads was used to set Definite Focus. 25 µL of septins at the desired concentration (in SSB either containing 300 mM KCl or 200 mM KCl for experiments with final KCl concentrations of 75 and 100 mM KCL or 50 mM, respectively) were mixed in the reaction chamber, focus was confirmed, then acquisition was started.

### Measuring protein adsorption on planar supported bilayers

The mean grey values of the images of the bilayers with and without protein were collected in FIJI. Background subtraction was performed for each timestep by subtracting the mean value before protein was added. The values of the replicates were averaged and fit with a Hill function where n is constrained to ≥1.

### Measuring protein adsorption on microsphere supported bilayers

To collect the protein intensity on single beads and exclude clumps of beads, a mask was applied to the 561 channel of the timeseries to collect the area integrated intensity of 488 signal where 561 signal was 1-2 µm2. For a few experiments, the first 3-4 timepoints had unusually high background signal; we suspect this signal was caused by concentrated, undiffused protein because in all cases, after the 4th-5th timepoint the intensity values dropped to expected levels and the adsorption curves matched experiments without this abnormality. For these experiments, all values from those abnormal timepoints were removed as they do not represent the kinetic trends observed in this system. The values of the replicates were averaged and fit with a Hill function where n is constrained to ≥1.

### Fluorescence Correlation Spectroscopy

Diffusion of SNAP 488-labeld septins in solution was measured using a Zeiss LSM 980 laser scanning confocal equipped with two Zeiss BiG.2 detector that contains two GaAsP photomultiplier tube (PMT) detectors for performing fluorescence correlation spectroscopy (FCS). Protein diluted in reaction buffer was added to a reaction chamber on a PEG-coated coverslip and incubated for 30 min before collecting measurements. Four random locations in the reaction chambers (20 µm above the coverslip) were sampled, collecting 20 5 s measurements at each location using a 40X water objective. Autocorrelation data and average count rates for each measurement were exported using the fcsfile package (version 2024.5.24, https://github.com/cgohlke/fcsfiles) in Python, and subsequently imported into MATLAB (version 2023b, Mathworks) for fitting to the 2D, single-component autocorrelation function

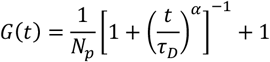

where *N*_*p*_ is the average number of labeled particles in the focal volume, *τ*_*D*_ is the diffusion time, and *α* is the anomalous diffusion coefficient. As a quality threshold, results were filtered by using the 95% CI of the fit parameters *N*_*p*_, *τ*_*D*_, and *α*, such that if the magnitudes of each parameter were greater than their respective 95% CI values, the measurement was not included in the data presented here. As a further threshold, measurements with average count rates less than 1 kHz were also excluded from analysis.

Counts per molecule (CPM) for each 5 s measurement was computed by dividing the average count rate by *N*_*p*_ and normalizing by the labeled percentage of the sample. The diffusion coefficient, *D*, was computed from *τ*_*D*_ using the relationship

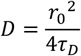

where *r*_0_ is the radius of the focal volume. *r*_0_ for our system was calculated by measuring *τ*_*D*_ of the fluorescent dye Atto 488, diluted in reaction buffer. Using the well-defined *D* of Atto 488, 400 µm^2^ s^-1^, we computed *r*_0_ using the above relationship.

### AlphaFold

The structure of the yeast septin octamer was predicted using the AlphaFold server. The query consisted of one copy of each of the 4 canonical yeast amino acid sequences. The prediction was run using standard settings. Models were visualized in ChimeraX and the unstructured regions of the CTEs and Cdc3 NTE were spatially adjusted for ease of viewing. Then using an AlphaFold predicted Cdc10-10 interface, the Cdc10 of two tetramers were matched to the 10-10 structure to assemble an octamer.

### Reactive Brownian dynamics simulations and analysis

#### Simulations

Simulations were performed using the molecular dynamics software HOOMD-blue version 2 (*24*) augmented with a custom C++ with python wrapper plugin, DyBond, which we adapted from a previous version first introduced as ‘epoxpy’ in (*39*), and which was shared with us by the authors (https://bitbucket.org/cmelab/hoomd_blue/src/dynamic_bonding/) via correspondence in the HOOMD-blue google group, ‘hoomd-users’.

2500 – 1000000 particles are initialized in a uniform cubic lattice in 3 dimensions in a box with periodic boundary conditions on all of the x-z and y-z planes, and with reflective walls at the top and bottom x-y planes with dimensions that specify particle concentrations used in the study (10^−9^ − 10^−5^ M). Reflective walls are implemented using the cut-off Lennard-Jones wall potential, 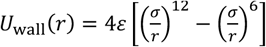, where *r* is the distance between the center of a particle and the wall, *ε* scales the strength of the potential, set to 5 for all simulations, *σ* is set to the radius of a particle, 0.5, and the potential is cut off after the distance is greater than 0.5, such that the potential is purely repulsive. Particles are initially equilibrated for 200 time steps, during which only repulsive wall interactions and excluded volume interactions among particles (described below) are active. Particles randomly update their positions according to Brownian motion, which is implemented using overdamped Langevin dynamics: ^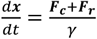^, where ***F***_***c***_ is all the forces acting on the particle with position ***x*** coming from all potentials and interactions, ***F***_***r***_ is a uniform random force, described next, and *γ* is the drag coefficient, set to 1 in all simulations. The random force has the following properties: ⟨***F***_***r***_⟩ = 0, ⟨|***F***_***r***_|^2^⟩ = 6*kTγ*/Δ*t*, where *kT* is the Boltzmann constant multiplied by the temperature, which is set to 1 in all simulations. Since the diffusion coefficient and the diameter of particles are both set to 1, a single simulation timestep corresponds to the average time it takes for a particle to diffuse a distance equal to its size.

After equilibration, particles are uniformly distributed throughout the simulation box. For simulations with a membrane interaction, an attractive cut-off Morse potential is activated at the bottom x-y plane of the simulation box: *U*_mem_(*r*) = *D*_0_[exp(−2*α*(*r* − *r*_0_)) − 2exp−(*α*(*r* − *r*_0_))], where *r* is the distance between the center of a particle and the membrane wall, *D*_0_ scales the strength of the potential and is set to 10 for simulations in the main text, and 5 (**Fig S1**) and 10 (**Fig S1**) for simulations in the supplement, *α* controls the range of the potential and is set to 2 in all simulations, *r*_0_ is the equilibrium distance between the membrane wall and the particle center and is set to 0.5 in all simulations, and the potential is only active at distances between 0.5 and 2.8. Simulations are integrated with Δ*t* = 2.5 × 10^−5^ for high concentrations, and Δ*t* = 5 × 10^−5^ for low and moderate concentrations. After equilibration in all simulations, pairwise bonding interactions among particles are activated, which we implemented using our DyBond plugin.

Our DyBond plugin adds several functionalities that were needed for this study to the standard molecular dynamics tools provided in HOOMD-blue described above. Specifically, it allows for the stochastic creation and breaking of pairwise bonds with specified rates between colliding and bound particles, respectively. During each update step, when the center-to-center distance of 2 unbound particles is less than or equal to the diameter of a particle (1), a new bond between them is created by generating a uniform random number between 0 and 1, and checking whether this number is lower than the annealing probability, which is set to 5 × 10^−3^ for simulations in the main text, and 5 × 10^−2^ (**Fig S1**) and 5 × 10^−4^ (**Fig S1**) for simulations in the supplement. The pairwise bonds are implemented using the harmonic potential, 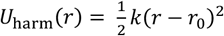, where *r* is the center-to-center distance between the 2 particles, *r*_0_ is the equilibrium rest length of the bond, set to the diameter of the particles in all simulations, and *k* scales the strength of the potential, set to 40 for all simulations. For all triplets of bound particles, DyBond adds an angle potential to enforce linear arrangements of bound particles into filaments. The angle potential we used is the cosine squared potential, 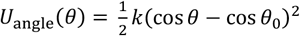, where *θ* is the angle formed by a triplet of particles, *θ*_0_ is the equilibrium rest angle, set to *π* for all simulations, and *k* scales the strength of the potential, set to 100 for all simulations. If the bond is not created, the excluded volume interaction will separate the particles, given by the cutoff harmonic potential, 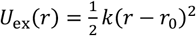, where *r* is the center-to-center distance between the 2 particles, *r*_0_ is the equilibrium rest length of the interaction, set to the diameter of the particles in all simulations, and *k* scales the strength of the potential, set to 50 for all simulations, and the potential is cut off at distances less than 0 and greater than 1, such that it only acts repulsively. Bonds are also stochastically removed during simulations. During each update step, each pairwise harmonic bond is removed if a uniform random number between 0 and 1 is less than the fragmenting probability, set to 3.94 × 10^−4^ in a single time step fo simulation time (i.e. 20000Δ*t* or 40000Δ*t*) for all simulations. When a bond is removed, all 3-particle angle bonds in which it participated are also removed.

Simulations were run until numbers of adsorbed particles, mean lengths, and fraction of aggregation values all reached steady state, which depended on concentration and whether a membrane interaction was present and ranged from hundreds to tens of thousands of simulation time steps.

#### Analysis

Output simulation .gsd files were analyzed by first doing a cluster analysis at each time step using the ovito python module. Particles are counted as a cluster if they are connected by bonds implemented by DyBond, such that clusters correspond to filaments. These cluster files are further analyzed using an R script, which calculates the fraction of aggregation (*φ*), defined as the fraction of particles in the system that are not monomers, and mean length at each time point for the bulk simulations. For the membrane simulations, these quantities are calculated only for those particles which are bound to the membrane, which we define as all particles within 1.5 distance units of the membrane. For the membrane simulations, we additionally calculate the membrane adsorption, defined as the total number of particles within 1.5 distance units of the membrane, at each time point. The *φ* vs time curves are first normalized by their steady-state values, and they are well fit by a single exponential function, *φ*_norm_(*t*) = 1 − exp(−*t*/*τ*), where *τ* is the timescale that is fit. The time at which *φ* reaches half of its maximum value, *t*_1/2_ *φ*, is given by log_2_ *τ*.

### Hill function fits

The FCS *τ*_*D*_ versus concentration data were first normalized to their maximum values at each salt concentration to facilitate fitting Hill functions. The following function was used to fit curves at each salt concentration: *τ*_*D*_(*c*) = (1 + (*K*_*a*_/*c*)^*n*^)^−1^ + *z*, where *cc* is the septin concentration, *K*_*a*_ is a fit parameter controlling the concentration at which 50% of the maximum *τ*_*D*_ is reached, *n* is the fit cooperativity parameter, and *z* is an offset. For all *τ*_*D*_ versus concentration data, *z* is set to the average *τ*_*D*_ of the non-polymerizable mutants at 2nM concentrations across all salt concentrations, since this is the lowest *τ*_*D*_ we would expect to measure with a non-zero septin concentration. The function was fit with the function ‘curve_fit’ from the python module scipy, with the constraints that *K*_*a*_ > 0 and *n* ≥ 1. The fit values of *n* are 1.62 ± 0.33 for the 50mM KCl data, 4.5 ± 0.64 for the 75mM KCl data, and 12.42 ± 3.78 for the 100mM KCl data, where the error is the square root of the diagonal elements in the covariance matrix output by ‘curve_fit’, which quantifies the uncertainty in the *n* parameter fit. The planar bilayer adsorption data were fit in an analogous way, except the offset *z* was set to 0 for all fits. The bead assay adsorption data were fit in an analogous way, except the offset *z* was set to the value of the data at time 0.

### Isodesmic model predictions for *τ*_*D*_ versus concentration observed with FCS

Our goal is to relate the average diffusion time of a population of septin octamers and filaments to the bulk concentration, since this is what is observed by FCS. Assuming isodesmic polymerization, we can derive an expression relating the average filament length to the bulk concentration of septins using the following results. Oosawa and Kasai (*40*) obtained the following relations for an isodesmic polymer that grows by monomer end-on addition:

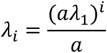

where *a* is the association constant, *λ*_*1*_ is the number concentration of monomers, and *λ*_*i*_ is the number concentration of *i*-mers; and

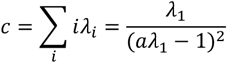

where *c* is the bulk concentration in the system. Septins undergo not only end-on monomer addition, but also coagulation and fragmentation. However, Markvoort *et al*. (*25*) have shown that the inclusion of coagulation and fragmentation reactions in an end-on monomer addition isodesmic model alters kinetics but retains the same steady-state properties. Therefore, the model from (*40*) is appropriate for hypothesizing about septins.

Since the diffusion time measured by FCS represents an average over the diffusion times of all species present at a given condition (i.e. bulk concentration) that happen to diffuse through the focal volume, we are interested in the weight-average length of the isodesmic polymer as a function of bulk concentration. The intuition is that a filament that is twice as long is twice as likely to be observed. Using the relations above, the weight-average length is:

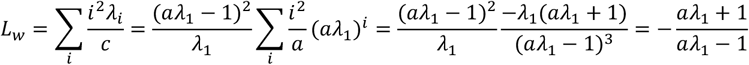

when |*aλ*_1_| < 1, which is always satisfied for reasons that will soon be clear. We now have an expression for the weight-average length of an isodesmic polymer as a function of the association constant and the number concentration of monomers in the system. We instead would like to relate the weight-average length to the bulk concentration, since this is the experimentally known value. Again, we can depend on the above relations:

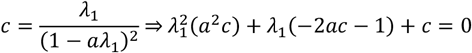

Using the quadratic formula, we obtain the desired result:

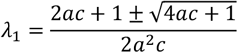

Here, we see that when 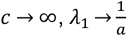, satisfying the inequality that is required above to simplify the summation in the expression for *L*_*w*_. Plugging in this expression for *λ*_*1*_ gives the dependence of the weight-average length on the bulk concentration and the association constant:

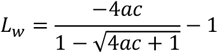

We next need to obtain the expected translational diffusion coefficient for freely rotating rigid rods in solution. From Doi (*41*), we have the following expression for a diffusing rod with an isotropic angular distribution:

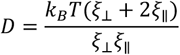

where *k*_*B*_ is Boltzmann’s constant, *T* is temperature, *ξ*_⊥_ is the friction coefficient perpendicular to the long axis of the rod, and *ξ*_∥_ is the friction coefficient parallel to the long axis of the rod. The diffusion coefficient can be related to the diffusion time measured by FCS by the following relation from Elson (*42*):

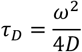

where *ω* is is radius of the focal volume. Thus, for a freely rotating rod, the diffusion time is given by:

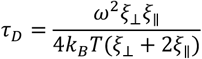

The expressions from (*43*) relate the friction coefficients to the dimensions of the rod:

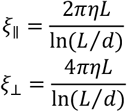

where *η* is the viscosity of the medium, *L* is the rod length, and *d* is the rod diameter. Plugging in these expressions, we obtain the diffusion time for a rod as a function of its length and diameter:

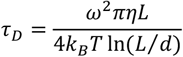

For septins, we set *d* = 4nm, and for *L* we plug in the weight-average length such that we have a relationship for the diffusion time as a function of the bulk concentration of septins in the system, assuming isodesmic polymerization. We see that the isodesmic model always requires a concave function relating bulk concentration to diffusion time. Such a shape always requires a Hill coefficient fit to be ≤ 1, indicating no cooperativity. However, in experiments we see that *τ*_*D*_ versus concentration curves are always well-fit by Hill coefficients greater than 1, indicating that polymerization in solution is not isodesmic.

## RESULTS

### A simulation to examine how binding to a surface contributes to cooperative polymerization

Given that septin structures are typically observed on membrane surfaces *in vivo*, it is plausible that this interaction could serve as a stabilizing or nucleating event in a nucleation-elongation scenario (*23*). Because septin assembly has appeared to be either isodesmic or cooperative based on context (*9, 13, 16*), we set out to examine to what extent a membrane interaction may induce features of cooperativity in an otherwise isodesmic polymer using coarse-grained simulations. We created a reactive Brownian dynamics model by adding chemical reactions with a specified number of bonds and reaction rates to the HOOMD-blue molecular dynamics software (*24*) using a custom plugin (see methods). We simulated individual septin octamers as spheres that can bind at most 2 additional partners as linear filaments in a 3D box with and without a membrane interaction at the bottom surface using overdamped Langevin dynamics (**Fig 1E**). Because the exact interfaces for lateral septin interactions remain unknown, we chose to exclude this feature from the model.

For simulations without a membrane interaction, we modeled polymerization using an isodesmic model with coagulation and fragmentation reactions: 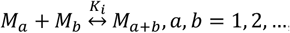, where *M*_*a*_ is a filament made up of *a* bonded monomers (**Movie S1-2**). Septins have been observed to anneal and fragment anywhere along their length (*9*), and isodesmic polymerization with such coagulation and fragmentation reactions have been shown to speed up relaxation to equilibrium, but not to alter the equilibrium state relative to an isodesmic model in which monomers can only exchange at the ends of filaments (*25*). We added a membrane interaction to a second set of simulations by introducing a binding energy with the bottom surface of the simulation box: 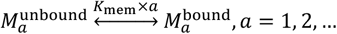, where *K*_mem_ is the binding energy of a single monomer to the membrane. This is implemented as a short-range interaction such that each septin monomer near the “membrane” feels an equivalent attraction to the surface (**Movie S1-2**). Septin filaments can still diffuse once they are bound to the simulated membrane surface.

For systems with and without membrane interactions, we simulated polymerization over 3.5 orders of magnitude of protein concentrations, starting from a diffuse gas of monomers until steady-state. We tracked the fraction of polymerization (aggregation) (φ) over time in the bulk or on the membrane, respectively, which is defined as the fraction of the system that is not monomeric (**Fig 1F**). We found that for both systems, kinetic curves never exhibited lag phases and were fit well with single exponentials, indicating a constantly decreasing slope with time that is characteristic of isodesmic assembly (**Fig 1F**). Similar kinetic curves of average length and total membrane adsorption over time show the same features (**Fig S1**). Plots of *tt*_1/2_ for φ versus concentration reveal linear trends in log-log space for both systems, also indicating isodesmic assembly (**Fig 1G**). By observing steady-state average lengths versus concentration for each system, we see that while membrane interactions indeed promote enhanced assembly and longer lengths, they do not result in a critical concentration as would be expected for a nucleation-elongation mechanism (**Fig 1H**). Plots of membrane adsorption over time also show constantly decreasing slopes over all tested concentrations (**Fig 1I**). These simulations show that recruitment to a 2D surface is not sufficient to introduce cooperative properties to an isodesmic polymer system.

It is possible that the apparent lack of cooperativity on a surface is due parameter choices in the simulation. We simulated additional systems with membrane interactions at two other parameter regimes corresponding to fast membrane binding and slow polymerization, and vice versa, and similarly found many signatures of isodesmic assembly but none corresponding to cooperative behavior (**Fig S1**). We therefore conclude that membrane binding does not necessarily induce cooperativity in an otherwise isodesmic polymer, contrary to a recent viewpoint (*23*). We expect this result, since membrane binding does not introduce a new, length-dependent association constant for polymerization. In principle, the membrane unbinding rate of filaments, *k*_off_, does scale with filament length. However, we expect the probability of a filament to unbind in time Δ*t* to immediately approach 0 relative to all other timescales upon filament assembly, since all monomers in a filament must simultaneously unbind for a filament to unbind, such that: 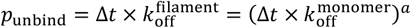, for a filament of length *a*. Therefore, membrane unbinding rates do not affect polymerization but only serve to contribute to the steady-state concentration of monomers on the membrane. Membrane binding does enhance isodesmic polymerization by restricting proteins to 2 dimensions, effectively re-scaling concentrations to higher values which promotes increased assembly. While septins often, if not exclusively, assemble on a substrate in cells such as a membrane or another cytoskeletal network, this simulation shows that simply polymerizing on a substrate is not sufficient to introduce cooperative features to the process. We thus set out here to characterize the mechanisms of septin assembly in different contexts and different ionic conditions.

### Septins have features of cooperative polymerization in solution

Although the mechanism of septin polymerization has not been systematically examined, previous observations have revealed trends consistent with both isodesmic and cooperative processes. Specifically, cooperative recruitment to curved membrane surfaces and isodesmic features on planar-supported membranes have both been observed (*9, 13, 16*). While the geometry of the membrane is clearly different in these previous studies, the buffer conditions of those two contexts were also different because of the different assembly kinetics on flat and curved surfaces and the need to maintain membrane integrity and limit photobleaching. Therefore, it is possible that the differences in mode of assembly are due to membrane geometry and/or ionic conditions. The effect of ionic strength on septin assembly has been examined at high protein concentrations in a wide range of KCl concentrations (*26, 27*). In the following experiments, we also vary ionic strength, probing septin polymerization at low protein concentrations, with an initial focus on assembly in solution at three KCl concentrations (50, 75, and 100 mM). Importantly, these KCl concentrations are based on previous work and represent a standard range for *in vitro* cytoskeleton polymerization assays (*28, 29*).

To assess polymerization in solution, we used fluorescence correlation spectroscopy (FCS) to measure the diffusion and brightness of recombinantly-expressed, fluorescently-labeled septin complexes to track their assembly into small filaments below the diffraction limit. We used a non-polymerizable mutant that we previously generated, cdc11α6 (11α6), to calibrate measurements, as it is the minimal 32 nm protomeric state (**Fig S2A-C)**. In all conditions, recombinant septin complexes were diluted in reaction buffer containing 50, 75, or 100 mM KCl, and loaded into a chamber assembled on a PEG-passivated coverslip. The FCS time constant, *τ*_*D*_, and the calculated diffusion coefficient, *D*, reflect the time molecules spend in the focal volume and is a readout of both the shape and size of septin assemblies (**Fig 2A, B**). Based on the average *τ*_*D*_ of the non-polymerizable mutant, 11α6 (0.49 ± 0.06 ms), we expect conditions with average *τ*_*D*_ under 0.55 ms (*D* ∼ 22 µm/s^2^) to represent mostly, if not entirely unpolymerized septin complexes. As expected, reducing salt concentrations lead to slower diffusing (i.e. longer) filaments at lower septin concentrations compared to high salt (**Fig 2A, B**). While all salt concentrations showed some degree of cooperativity, there was a clear effect of higher salt increasing cooperativity (**Fig 2A**). Fits of the Hill coefficient, *n*, were 1.62 ± 0.33 for the 50mM KCl data, 4.5 ± 0.64 for the 75mM KCl data, and 12.42 ± 3.78 for the 100mM KCl data, in which values larger than 1 indicate positive cooperativity and the errors are standard error of the mean of parameter estimate uncertainties. A caveat is that the *n* estimate for 100 mM KCl is uncertain, as the data capture only a small amount of the curve. We derived the expected relationship between *τ*_*D*_ and concentration for filaments that undergo isodesmic polymerization and found that such curves will always be concave, leading to fits of the Hill coefficient that are ≤ 1. We therefore conclude that our data indicates that septin polymerization is not isodesmic in solution. Although there is a limited ability to detect an apparent critical concentration at 50 mM KCl, we observe an apparent critical concentration of approximately 3 nM at 75 mM KCl, and a higher critical concentration of approximately 12 nM at 100 mM KCl. Notably, these critical concentration estimates are at least an order of magnitude less than the estimated cytosolic concentration of septins in yeast where polymerized septins were not detectable in the cytosol, potentially due to the higher ionic conditions of the cytosol (*9*).

**Figure 2:**
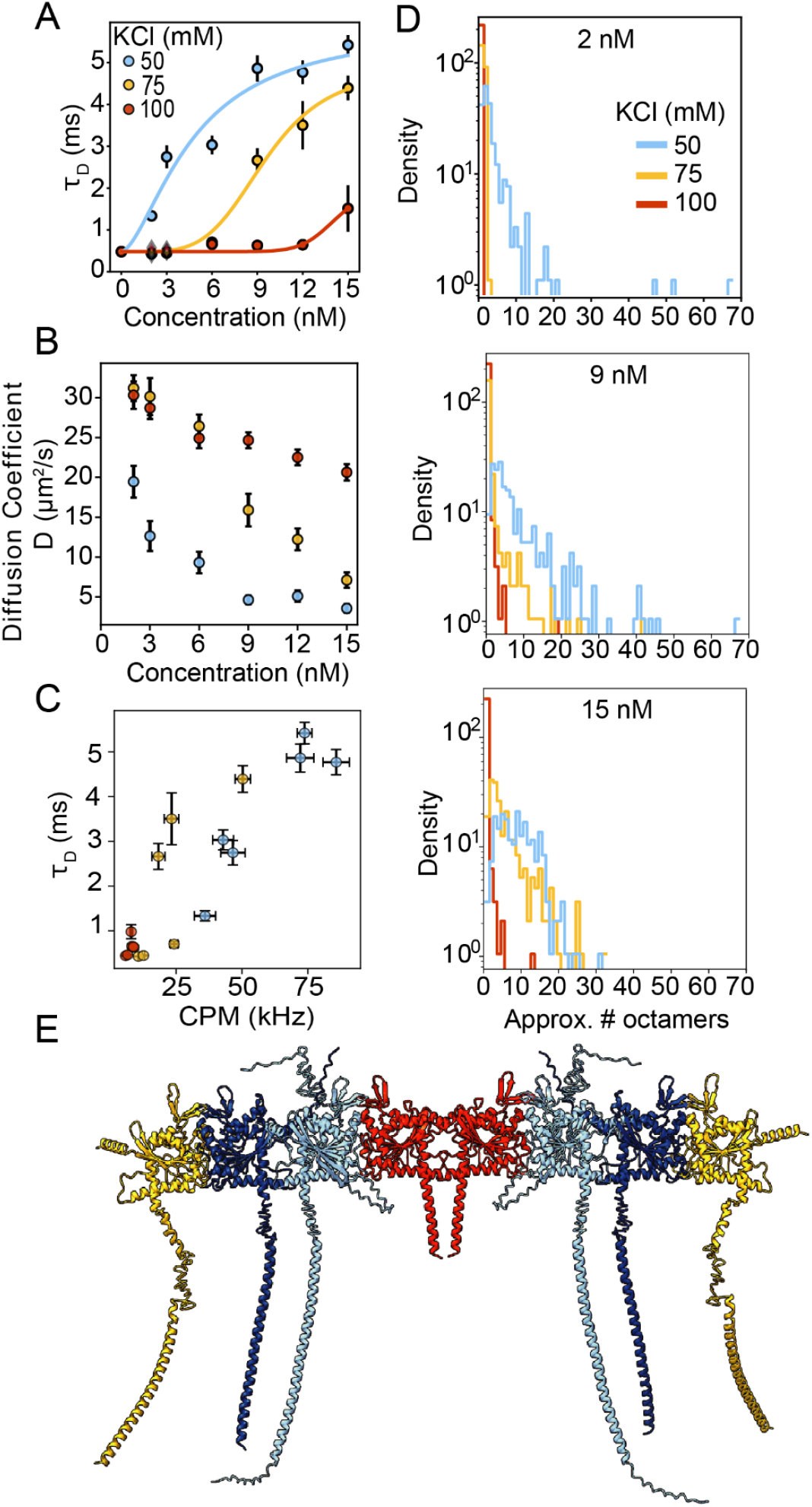
Septin polymerization and geometry in solution are impacted by salt. FCS determined diffusion time constants (A), diffusion coefficients (B) and counts per molecule of recombinant septin complexes at low salt (50 mM KCl, blue), medium salt (75 mM KCl, yellow), and high salt (100 mM KCl, red). A) *τ*_*D*_, the time constant for diffusion, plotted as circles for WT complexes and diamonds for non-polymerizable 11α6 complexes. These data were fit with Hill functions with the cooperativity parameter, *n* constrained to be ≥1, and the offset set to the average *τ*_*D*_ of 11α6 data at 2 nM across all salts. At low salt, *n* = 1.62 ± 0.33, at medium salt, *n* = 4.5 ± 0.64, and at high salt, *n* = 12.42 ± 3.78. B) Diffusion coefficients of septins. C) Counts per molecule (CPM) vs *τ*_*D*_. Error bars in A-C represent 95 % CI, error in Hill coefficients is SEM. D) Filament length distributions, estimated from CPM values, at 2, 9, and 15 nM protein. E) AlphaFold predicted structure of yeast septin hetero-octamer.

We next examined the relationship between *τ*_*D*_ and counts per molecule (CPM), which reflects the number of labeled subunits per complex and thereby serves as a proxy for mass (**Fig 2C**). Interestingly, we found that for a given mass (CPM), septins display different *τ*_*D*_ values in a salt-dependent manner, meaning that complexes of a given mass can adopt distinct geometries depending on salt. Specifically, complexes of a fixed mass appeared more compact as salt concentration decreased from 75 to 50 mM. Also, for a given salt, *τ*_*D*_ appeared to initially grow slowly with CPM, then super-linearly before eventually saturating (**Fig 2C**). This behavior indicates that there is likely some change in geometry of assemblies as septins polymerize in solution at higher concentrations, potentially corresponding to a relatively disordered nucleus that transitions into linear filament growth. The possibility of a large extended conformational state is supported by the predicted extended C-terminal ends of Cdc3, 11, and 12, and the N-terminus of Cdc3, which contain disordered sequences capable of adopting different conformations upon oligomerizing or bundling (**Fig 2E**). After nucleation, complexes may grow through both linear, end-on annealing, which lengthens filaments, and subsequent lateral adhesion or bundling of existing filaments, which thickens them. These various shapes (compact/disordered, linear, bundled) potentially explain the observed trends in **Fig 2C**.

Finally, we analyzed the length distribution of these filaments by plotting the number of octamers estimated in a complex based on CPM measurements (**Fig 2D**). As salt decreased, we observed a concentration-dependent decrease in *D* (**Fig 2B**) and a broadening of filament length distributions (**Fig 2D**). For example, complexes were predominantly small, under five octamers, at 75 mM KCl and 2 nM septins (**Fig 2D**). This trend is shown across all tested conditions in **Fig S2D**. At the same salt concentration, *D* decreased and filament length distributions shifted to higher values as septin concentration increased to 9 and 15 nM (**Fig 2B, D**). Of note, septins formed filaments in 50 mM KCl at the lowest measured septin concentration of 2 nM (**Fig 2D**). Finally, at 50 mM KCl and 15 nM septins, we observed a peak in the distribution rather than exponential decay (**Fig 2D**), which indicates non-isodesmic polymerization in solution and is consistent with the fit Hill coefficients that suggest cooperative assembly (**Fig 2A**).

### Ionic strength tunes the mode of septin assembly on supported bilayers

We next evaluated ionic effects on the septin assembly process on planar membranes (75% DOPC, 25% PI, 0.05% Rh-PE) with TIRF microscopy (**Fig 3**). Here, experiments were performed at 50 and 75 mM but not at 100 mM KCl, as assembly was too slow to reliably measure (**Fig S3**). Salt concentration dramatically changed the rate of filament elongation and appeared to impact the apparent mode of polymerization (**Fig 3A, B**). Consistent with simulations in **Fig 1**, the planar membrane did not itself enhance cooperativity that we detected in solution. Specifically, the time courses in **Fig 3A** show that septin assembly on membranes was either very weakly cooperative or not cooperative at 50 mM KCl, but transitioned to clearly cooperative, albeit with a modest lag time, at 75 mM KCl (**Fig 3A**). At 50 mM KCl, septin adsorption was rapid, and filaments above the diffraction limit formed within 10 min of addition (**Fig 3A**). Because there was no lag phase at 50 mM KCl, with a low Hill coefficient near 1, polymerization was inconsistent with a cooperative system. There are two possible explanations for this: 1) Membrane binding competes for an additional binding site that is otherwise available in solution, thereby eliminating a possible cooperative effect, or 2) Assembly is so favorable at 50mM KCl that nucleus formation and subsequent elongation are so rapid (“downhill nucleation”) that we fail to detect the regime corresponding to the weaker nucleus association constant, *K*_*a*_. However, the 75 mM KCl data appears cooperative, similar to the corresponding

**Figure 3:**
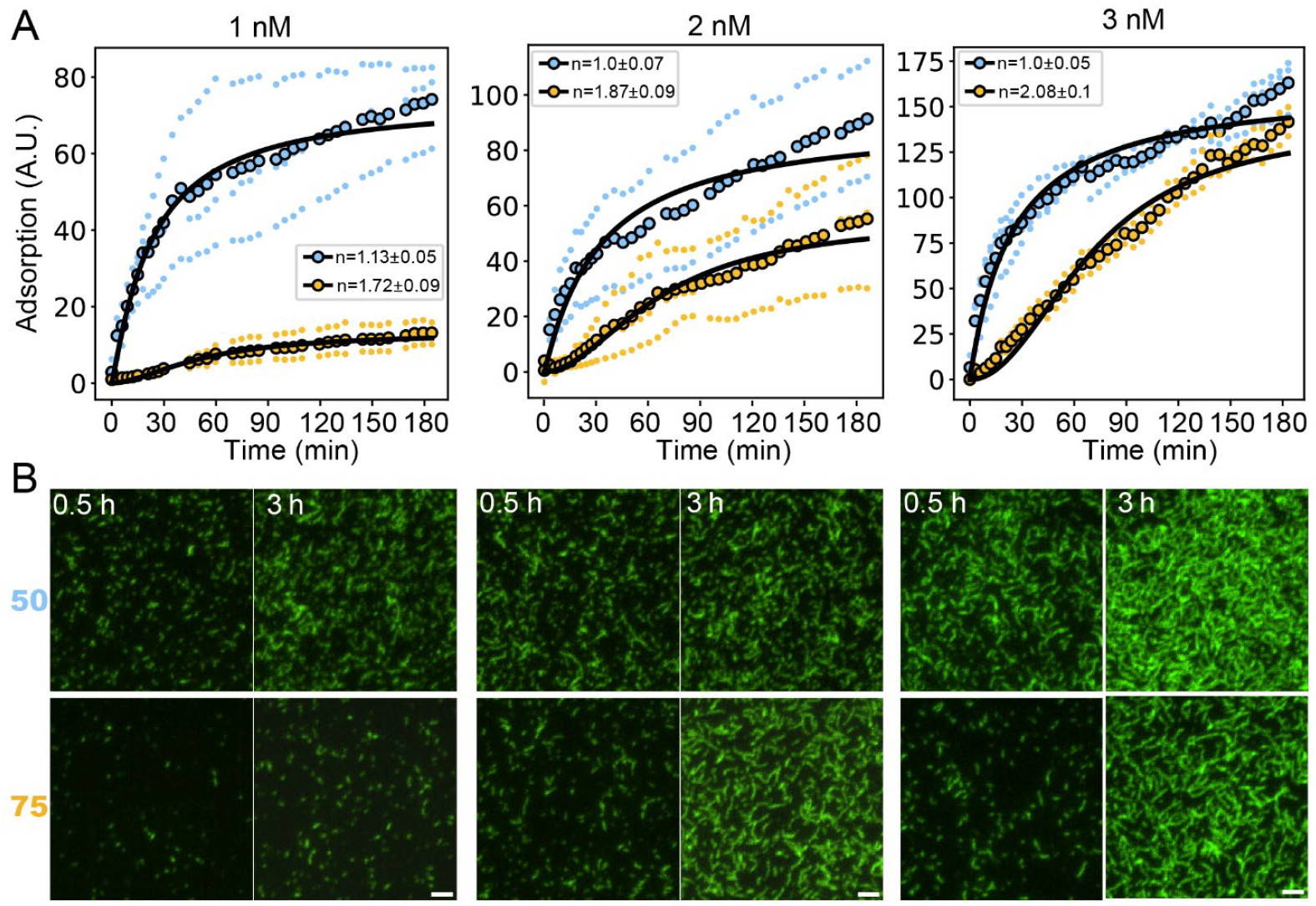
Higher ionic strength reduces septin polymerization and shifts polymerization modes on planar membranes. A) Quantification of septin adsorption on planar SLBs. Distributions were fit with the Hill function where *n* is constrained to be ≥1 (blue line, 50 mM KCl, yellow line, 75 mM KCl). B) Representative TIRF images of polymerized septin complexes after 0.5 and 3 h on planar SLB (75% DOPC, 25% PI, 0.05% Rh-PE). Images represent the plotted data above in A)- right 1 nM, middle 2 nM, and left 3 nM. All micrographs scaled and contrasted equally, scale bar, 2 µm.

FCS data (**Fig 2A**) albeit with weaker signatures of cooperativity (**Fig 3A**). It is possible that some or all of the same effects hypothesized above for 50 mM are at play at 75 mM KCl. Notably, at 75 mM KCl, in addition to slow membrane adsorption and filament formation, a substantial portion of the protein remained in solution compared to 50 mM KCl based on their saturation values. The high soluble pool may have been due to charge screening, as higher ionic strength could limit electrostatic-based recruitment of protein to the membrane. Another possible explanation that is not mutually exclusive is that slower elongation rates at 75 mM KCl produced shorter filaments with higher dissociation rates from the membrane. In conclusion, the presence of an attractive membrane did not enhance cooperative modes of polymerization, instead minimizing the role of cooperativity potentially due to a reduction in the conformational flexibility of septins when bound to a membrane.

### Cooperative adsorption on curved geometries varies in different salt regimes

In many cell contexts, septins are preferentially enriched at locations with micron-scale membrane curvature. Our previous work with both reconstitution experiments (*13, 14*) and physical modeling (*16*) of these curved assemblies suggested that there is a cooperative component to curvature-dependent adsorption with multiple distinct equilibria due to features such as filament packing and layering. To measure salt-dependent septin adsorption over time on curved membranes, we coated 1 µm glass beads with SLBs comprising the same lipid mixtures used in planar membrane experiments but supplemented with 2% Biotin-PE, which enabled tethering of beads to Biotin-PEG-conjugated coverslips via NeutrAvidin to facilitate imaging with confocal microscopy (**Fig 4**). In 50 mM KCl, septins rapidly adsorbed to curved membranes, reaching saturation within 0.5 h (**Fig 4A**). In 75 mM KCl, septins adsorbed more slowly, reaching saturation within approximately 3 h (**Fig 4A**). Notably, membrane curvature appeared to enhance adsorption at all KCl concentrations, as we were able to observe assembly at 100 mM KCl that was not evident in the planar system. At 100 mM KCl, septin accumulation on the membrane seemed to follow multiple regimes, including a lag and plateau phase within 1 h followed by an additional adsorption phase until reaching a second plateau within approximately 2.5 h (**Fig 4A**). These sequential steps in adsorption may arise from septin layering, with lower septin-septin affinity compared to septin-membrane affinity.

**Figure 4:**
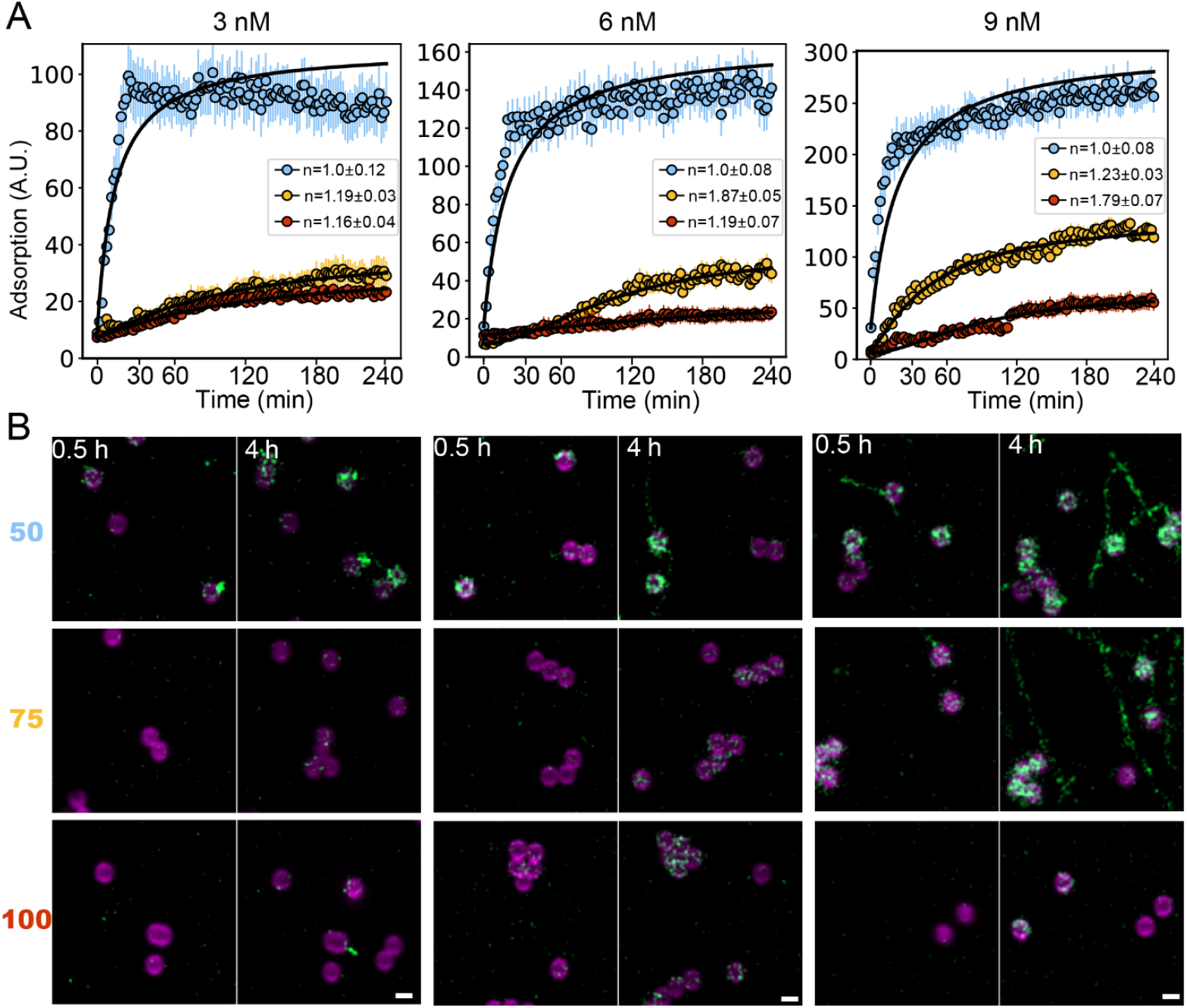
Septin assembly on curved membranes is enhanced at low salt. A) Quantification of septin adsorption onto individual 1 µm SLB-coated beads at low (50 mM KCl, blue), medium (75 mM KCl, yellow), and high (100 mM KCl, red) salt conditions. Distributions were fit with the Hill function, where *n* is constrained to be ≥1. Beads were imaged every 2 minutes for 4 hours. Data are represented as the mean ± 95% Cl. N > 500 beads for each condition, from 3 different experiments. B) Representative Airyscan images of SLB-coated 1 µm beads (73% DOPC, 25% PI, 2% Biotin-PE, 0.05% Rh-PE, magenta) after 0.5 and 4 h with septin complexes (green). Top-50 mM KCl, Middle- 75 mM KCl, Bottom-100 mM KCl. Images contrasted equally within each protein concentration, all scaled equally. Scale bar, 1 µm.

Analysis of cooperativity on curved membranes mirrored our findings from planar membranes. Specifically, we again saw that 50 mM KCl followed a non-cooperative mechanism based on fit Hill coefficients (**Fig 4A**), potentially owing to the two scenarios suggested above for planar membranes: 1) a brief, undetectable lag period and/or 2) conformational changes that occlude binding sites when membrane-bound. At higher KCl concentrations, weak cooperativity became apparent that may have been even weaker compared to planar membranes (**Fig 4A**). This weakening of cooperativity could be explained by more energetically favorable nucleation and elongation on curved membranes, which may have reduced lag times and/or induced structural changes that hindered cooperativity and promoted more rapid assembly. In all conditions, bead coverage was heterogeneous: some beads were populated early and became saturated by the end of the time course, whereas a small fraction were not bound at all (**Fig S4**). Importantly, this behavior is expected for a cooperative system, which may indicate that lower salt concentrations lead to nucleation timescales faster than our detection limit.

Because the data shown here are restricted to single beads, plots in **Fig 4A** do not capture the presence and growth of filaments in solution at lower salt concentrations as expected from our FCS measurements (**Fig 2**). However, images do reveal solution filaments at every tested septin concentration in 50 mM KCl (**Fig 4B**). Such filaments were visible as early as 20 min for 9 nM septins (**Fig 4B**). In 75 mM KCl, filaments were consistently apparent with 9 nM septins after 50 min (**Fig 4B**). Together, these data indicate that 1) septin adsorption is increased at curved membranes, 2) ionic effects on membrane adsorption and assembly are consistent between curved and planar membranes, and 3) septins preferentially assemble on membranes with lower apparent cooperativity compared to assembly in solution.

## DISCUSSION

Analysis of the mechanisms of polymerization has provided a critical framework for understanding the assembly and functions of the cytoskeleton. Cooperative polymerization mechanisms in actin and tubulin enable the cell to control precisely where and when polymers form through the action of regulatory nucleation proteins such as formins or gamma-tubulin (*44-45*). In this study we sought to understand the intrinsic polymerization modes of septins to ultimately build a framework for understanding how cells control their assembly in time and space. We find that septins display hallmarks of cooperative assembly that depend on salt and the geometry of membranes to which they bind.

Septin polymerization and substrate engagement depend strongly on electrostatic interactions. For example, membrane binding and polymerization require negatively charged lipids including phosphatidylinositol (PI), phosphatidylinositol-4,5-bisphosphate (PIP_2_), or phosphatidylserine (PS) (*9, 15, 30, 31*). Moreover, the earliest purifications and analyses of septin polymerization showed high sensitivity to buffer ionic strength, with less than 150 mM KCl generally required for polymerization in solution (*10-12*). The salt-sensitivity of filament formation via the Cdc11-11 interface is well-documented (*26*), but the direct impacts of ionic strength on polymerization have not been mechanistically dissected. Here we find remarkable salt-sensitivity in the polymerization mode (cooperative vs isodesmic), assembly rate, critical concentration, and membrane adsorption. Although the salt concentrations sampled here are below physiological levels, these regimes are regularly employed in assays of cytoskeletal polymerization. For example, the G-buffer used for measuring F-actin formation typically contains 50 mM KCl (*29*). By sampling a relatively narrow range of ionic conditions, we uncover how modest salt changes can have dramatic consequences on septin assembly. Such plasticity may be accessed in cells by the action of septin regulatory proteins, post-translational modifications, and/or changes to membrane lipid composition and geometry.

Septins exhibit apparent cooperative polymerization in solution at all salt concentrations tested. The degree of cooperativity and critical concentrations for polymer formation both increase with increasing salt. No study to date has found evidence of septins forming filaments in cells in the cytosol; this may be the result of the high ionic strength of cytosol, the presence of some kind of capping cofactor, and/or the need for a nucleation factor to overcome a low nucleation rate. In any case, our measurements show that at a range of salt and protein concentrations, septin filaments show hallmarks of cooperative assembly. This signature could suggest some conformational change induced in the protein complex upon oligomerization, favoring elongation; such a change would need to be allosteric in some way and propagate across the 32 nm complex to the free end as a site of elongation. There are hints of such long-range propagation of end-to-end information in septins during the formation of the oligomers such that all complexes have symmetric ends bearing Cdc11 or Shs1, but there are no mixed oligomers (*32*).

Our results suggest that septin filaments in solution follow distinct assembly regimes, growing isotropically at first, then anisotropically, and finally isotropically again as mass increases (**Fig 2C**). We suggest that these changes reflect different modes of growth including elongation and bundling, which has been well documented for animal septins in solution under certain circumstances (*33, 34*).

When tagged endogenously, septins have been nearly exclusively found assembling on membranes or cytoskeletal surfaces in cells. Assembly on a surface lowers critical concentrations and increases the rate of assembly by concentrating septins on 2 dimensions rather than 3. We first explored the consequences of membrane binding using simulations of an isodesmic polymer in solution and in the presence of an attractive membrane. We found no regime where membrane binding induced hallmarks of cooperative assembly, but we did find that it can accelerate the rate of filament elongation. Notably, when we experimentally measured adsorption on planar membranes, we saw a weakening of cooperative signatures at intermediate salt concentrations, and apparently isodesmic assembly at low salt concentrations. An apparent loss of cooperativity could be due to competition of the membrane with septin binding surfaces that promote cooperativity in solution, or because of “downhill nucleation” in which nucleus formation is energetically favorable and subsequent elongation is so rapid that we never observe the slower growth phase corresponding to nucleus formation. Additionally, it is possible that the elongation phase on membranes operates via an isodesmic mechanism (i.e. there is only 1 association constant, *K*_*e*_, for coagulation and fragmentation reactions after nucleus formation). Curved membranes appear to promote enhanced adsorption relative to planar systems since even high salt can support detectable adsorption. Based on fit Hill coefficients, curved membrane binding may further weaken cooperative signatures beyond what is observed for planar systems. Interpreting the nature and source of this cooperativity is challenging as there are many possible binding equilibria in a single adsorption curve including binding and elongation of filaments on the membrane, lateral interactions, and stacking of filaments. However, one possibility is that enhanced interactions between curved membranes and the amphipathic helices of Cdc12 (*14*) may further restrict the conformational freedom of growing filaments, occluding other possible septin-septin binding interfaces that may contribute to cooperativity in solution.

While no biochemically-characterized nucleators have been described for septins, it is known that assembly can be promoted by recruitment through Rho-GTPase signaling and effectors (*35*). Furthermore, enrichment of PI(4,5)P_2_ at the membrane, specifically at a preemptive bud site may contribute to septin recruitment and structural maintenance (*36, 37*). The combination of proper recruitment and concentration of septins to the membrane likely allows the proteins to overcome the ionic barrier of the cytosol and polymerize as needed. Together our findings demonstrate septin polymerization defies canonical cooperativity found in other cytoskeletal polymers.

## Author Contributions

EJDV performed the in vitro experiments, IS performed the simulations. EJDV, IS, and WTS analyzed the data. EJDV, IS, WTS, and ASG wrote the paper.

## Acknowledgements

The authors would like to thank Benjamin Woods for his work in nucleating this study. We thank Kevin Cannon for his feedback on this manuscript. This work was supported by NSF grant MCB-2016022 (ASG and EV), NIH T32 grant 5T32GM119999 (EV), NIH Pathway to Independence Award K99GM149757 (WTS), NIH T32 grant 2T32GM008570-21A1, the Max Planck Society, and the Alexander von Humboldt Foundation (IS).

