## SupplementalFigures for "Cooperativity in septin polymerization is tunable by ionic strength and membrane adsorption"

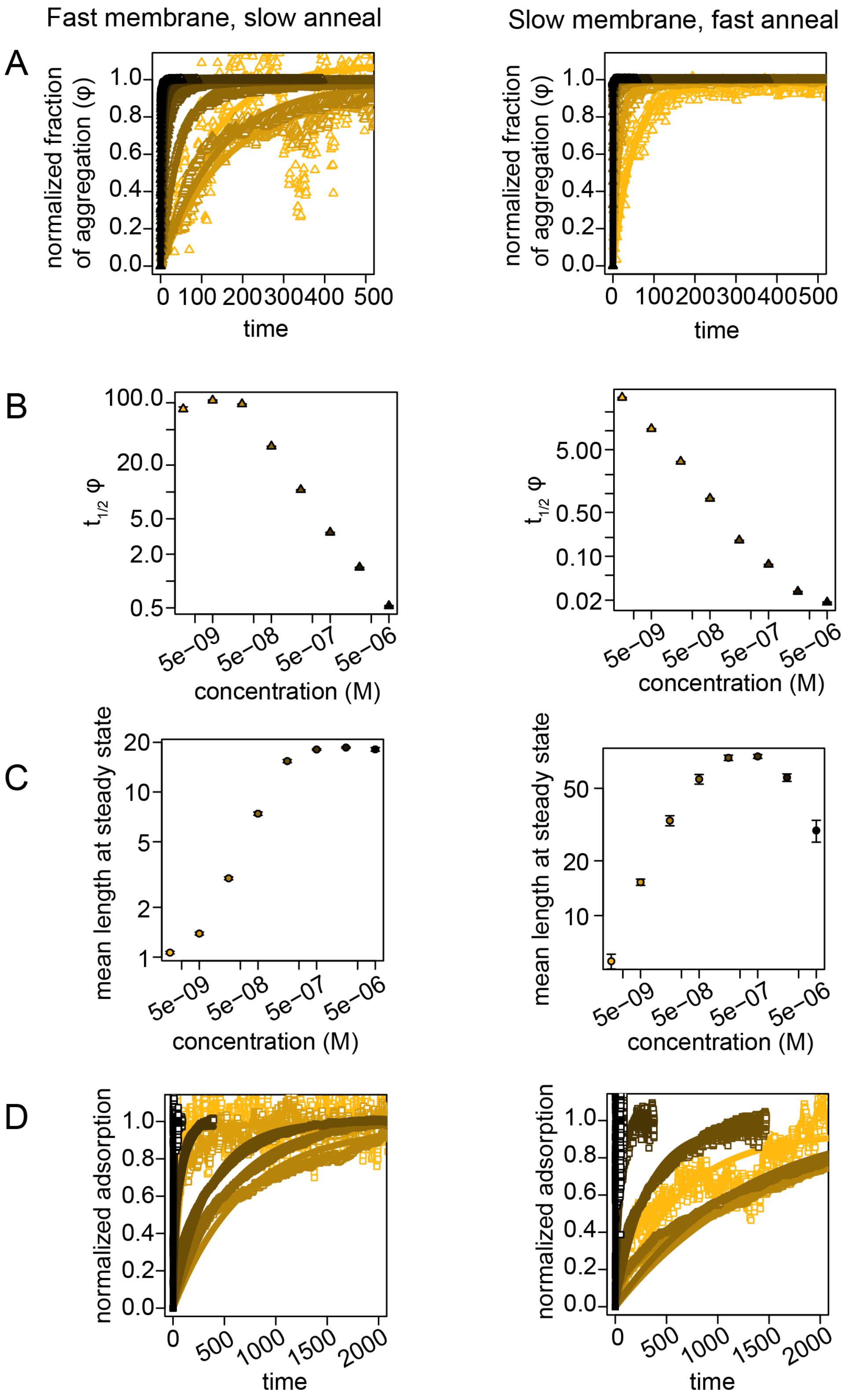


Figure S1- Simulating fast and slow adsorption and annealing also show membrane binding is not sufficient to induce cooperative assembly.

Left, $D_0$ , strength of the membrane potential set to 10 and annealing probability is set to $5×10^{-2}$
Right, $D_0$ set to 10 and annealing probability is set to $5×10^{-4}$

A)Fraction of aggregation over time at multiple concentrations  for simulations. B) Mean filament lengths at steady state across a range of concentrations for simulations with membranes. C)$\mathrm{Log}\, t_{1/2}$ for fraction of aggregation ($\varphi$) as a function of concentration for simulations. D) Simulated adsorption to the membrane over time.

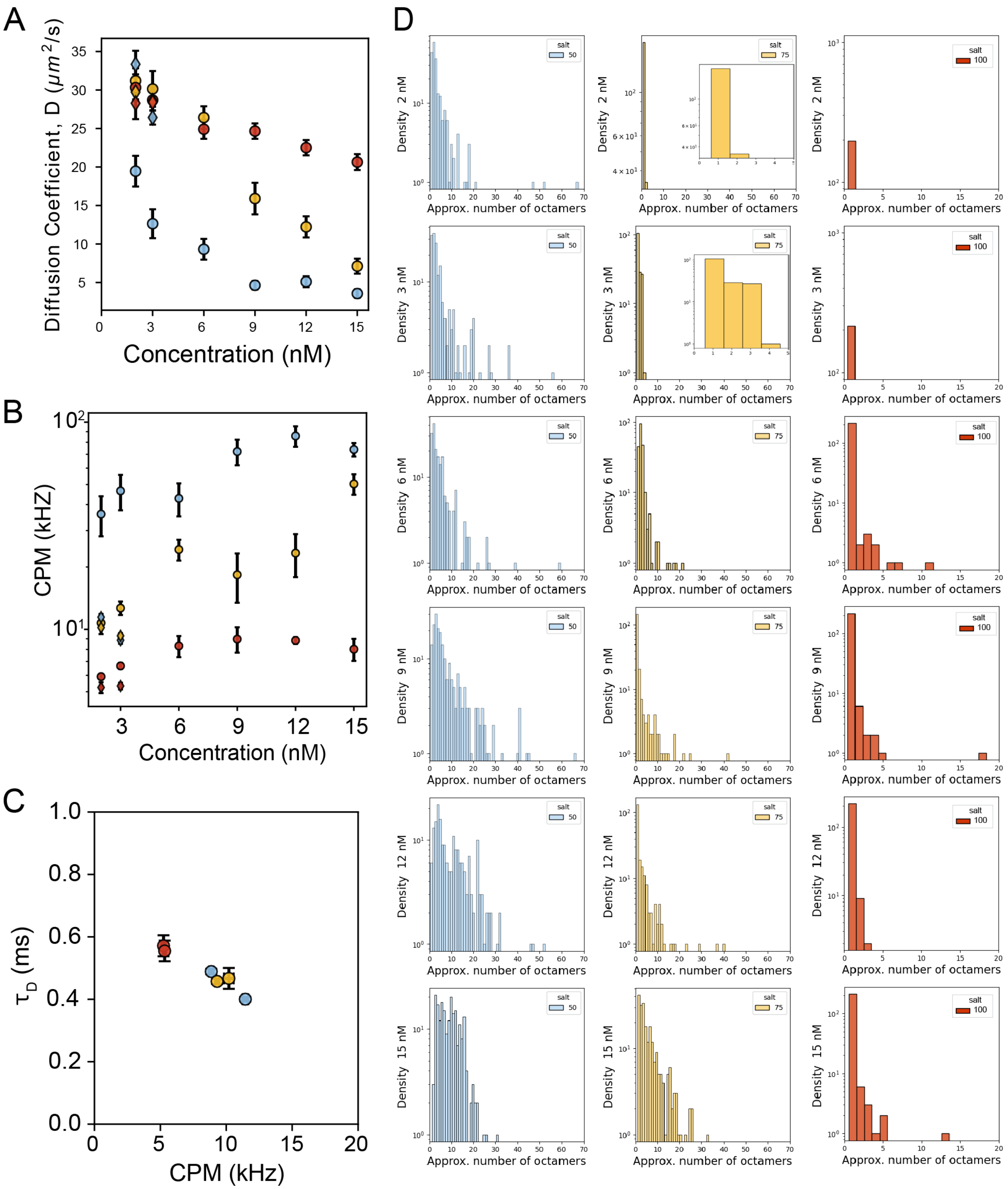


Figure S2

Diffusion coefficients (A) and counts per molecule (CPM) (B) of WT and 11α6 complexes plotted together, D) Counts per molecule (CPM) vs $\tau_D$ of 11α6 complexes. D) Filament length distributions. Error bars in A-C represent 95 % CI.

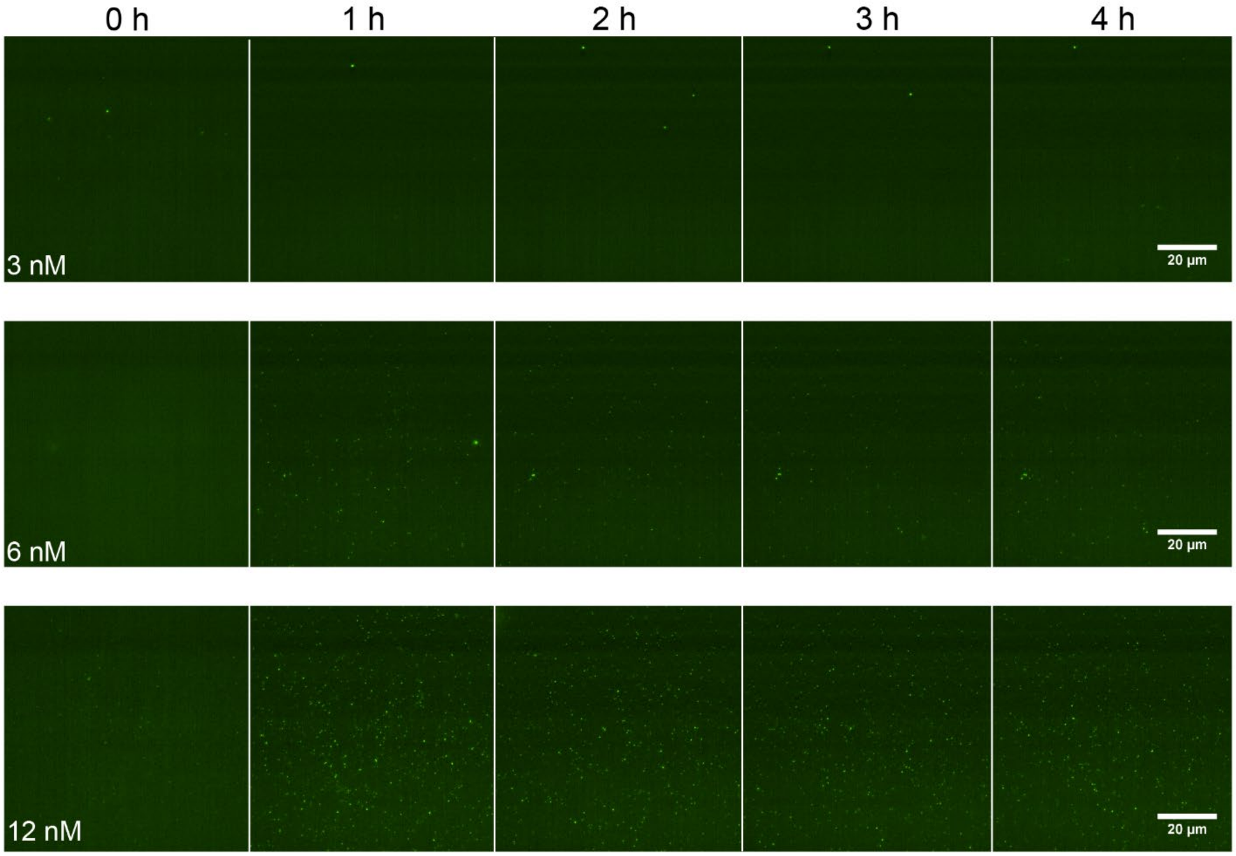


Figure S3

Whole field of view TIRF images of WT septin complexes planar SLB (75% DOPC, 25% PI, 0.05% Rh-PE) in 100 mM KCl buffer at 3, 6, and 12 nM.

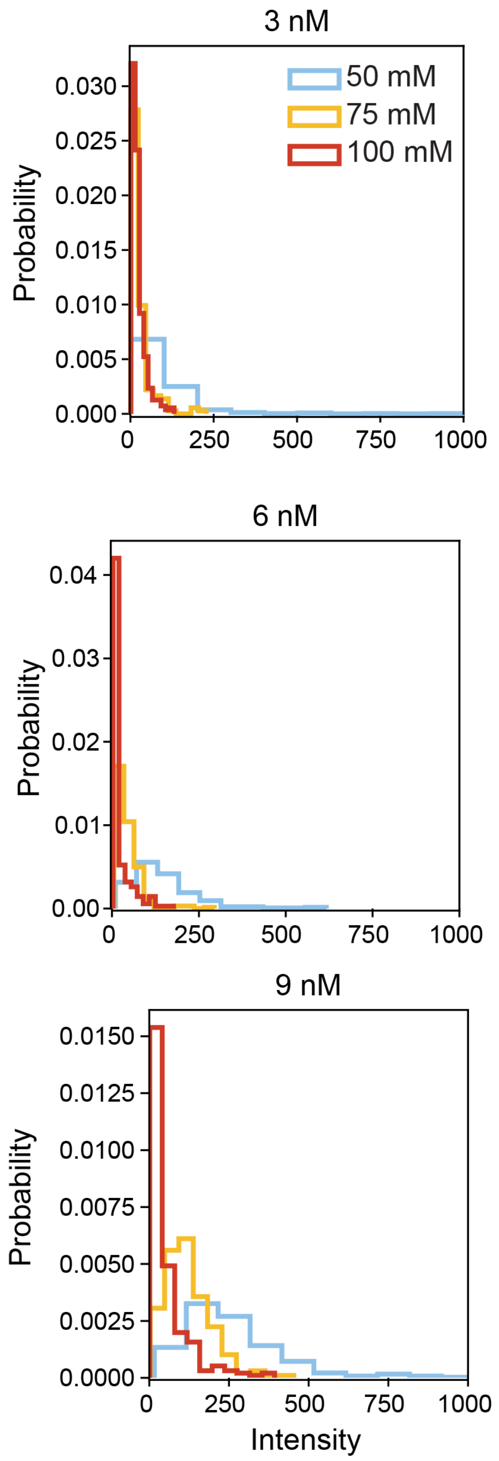


Figure S4

Probability distributions of protein intensity on onto individual 1 µm SLB-coated beads (73% DOPC, 25% PI, 2% Biotin-PE, 0.05% Rh-PE) at the final timepoint, t= 4h. N ≥ 120 beads

Movie S1

Simulation of bulk septin interactions without a membrane. Concentration is 1e-7 M, and the frame rate is 1 frame every 20 simulation time units. The colors appear when particles start to cluster after initial equilibration, with each color denoting a cluster.

Movie S2

Simulation of septin interactions in the presence of a membrane. Concentration is 1e-7 M, and the frame rate is 1 frame every 200 simulation time units. The colors appear when particles start to cluster after initial equilibration, with each color denoting a cluster.
